# Optogenetic stimulation of a cortical biohybrid implant guides goal directed behavior

**DOI:** 10.1101/2024.11.22.624907

**Authors:** Jennifer Brown, Kara M. Zappitelli, Paul M. Dawson, Eugene Yoon, Seton A. Schiraga, Amy E. Rochford, Mohamed Eltaeb, Arturo Rodriguez, Yifan Kong, Max Hodak, Alan R. Mardinly

## Abstract

Brain-computer interfaces (BCIs) hold exciting therapeutic potential, but tissue damage caused by probe insertion limits channel count. Biohybrid devices, in which the cell-device interface is crafted in the laboratory, hold promise to address this limitation, but these devices have lacked a demonstration of their applicability for BCI. We developed a biohybrid approach to engraft optogenetically-enabled neurons on the cortical surface housed in a 2D-scaffold of circular microwells. The engrafted neurons survived, exhibited spontaneous activity, and integrated with the host brain several weeks after implantation. We then trained mice with biohybrid implants to perform an optical stimulation task and showed that they could effectively report optogenetic stimulation of their neural graft. This demonstration shows that a cortical biohybrid implant can be used to transmit information to the brain of an implanted animal.

## Introduction

Brain-computer interfaces (BCIs) have shown promise for numerous clinical applications^1^, but their efficacy is limited by the small number of communication channels with the brain^2,3^. The density of neural recording channels is constrained by a number of factors, including brain damage caused by insertion of neural probes into the parenchyma. Tissue destruction following probe insertion causes an inflammatory response characterized by hypoxic injury, mobilization of microglia around the insertion site, and formation of a glial scar. These reactions result in neuron loss in the vicinity of recording probes, as well as vascular and glial damage that limits the duration that chronic implants can successfully record from neurons^4–9^.

Biohybrid neural interfaces are an emerging class of technology designed to circumvent these limitations by growing cells in a device in a laboratory and creating the device-host interface through a biological substrate^10,11^. Biohybrid devices are implanted into the target tissue in a manner allowing the implanted cells to engraft with host tissue and the embedded electrodes to record from and stimulate the engrafted cells. This approach holds potential advantages over traditional chronic neural probes; by allowing the engrafted cells to grow into the target tissue, rather than penetrating the tissue with an electrode, probes can be arrayed at potentially far higher density than otherwise possible. Neuron grafts delivered via microinjection are well known to integrate with the brain and acquire functional properties similar to the surrounding cortex, in some cases with therapeutic benefit^12–17^. However, neuron engraftment using microinjection has similar drawbacks to inserting neural probes: insertion of injection needles causes acute neuroinflammation and cell death early after engraftment^18–21^. Cortical biohybrid interfaces have been attempted using several different architectures, including seeding neural progenitor cells on silicon probes^22,23^, growing in projections from cells located distal to the cortex^24–27^, or by transplanting whole organoids and using an optical interface^28–31^. Biohybrid interfaces have also been implanted in the peripheral nervous system, including for regeneration of the neuromuscular junction^32–34^ and peripheral nerve fibers^35,36^. While biohybrid neural implants have exciting potential for BCI applications, they have so far lacked a functional proof-of-concept that they can be used to interface with the brain. A biohybrid implant used for this purpose would have to overcome three challenging problems: 1) the implant must have an architecture that could allow accessing a very large number of neurons at high resolution. 2) The transplanted neurons must exhibit a high rate of survival after engraftment. 3) The grafted neurons must functionally integrate with the brain in a manner that allows the implanted organism to access and use information from the graft.

To assess the suitability of this class of implants for BCIs, we designed a cortical biohybrid implant based on a planar array of microwells^37,38^ designed to hold one neuron per well. We implanted the biohybrid implant on the surface of the cortex, and observed robust survival and graft integration within several weeks after implantation. To assess functional engraftment into the brain, we developed an optogenetic stimulation task^39–41^ and tested whether mice could behaviorally report graft stimulation. Mice were able to perform this task to obtain rewards and sustain a positive bit rate. These results provide a proof-of-concept demonstration that a cortical biohybrid interface can provide input sufficient to drive goal-directed behavior, and opens a path for development of a class of high-bandwidth biohybrid neural interfaces.

## Results

To test the ability of a biohybrid implant to drive behavior, we designed an implant inspired by chronic mouse two-photon (2P) imaging, in which a piece of the skull is replaced by a glass coverslip^42^. We fabricated microwells designed to house single neurons and bonded them to the bottom of the coverslip so that the neurons loaded in the microwells could make contact with the brain, while also remaining optically accessible for light stimulation delivered through the coverslip. The microwell scaffolds were fabricated on fused silica wafers with SU-8 photoresist using photolithography. They were subsequently diced into 5×5mm dies, and then bonded to a glass coverslip with diameter 7 mm (Figure 1a). The microwells in the scaffold are packed in a hexagonal lattice at a 15 µm pitch. The interior of each microwell is a circle with a 10 µm diameter; the side walls are 2.5 µm thick and 8.6 µm long and each microwell is 15 µm deep (Figure 1b). This configuration allowed dense packing of cells in the scaffolds (approximately 1.18 x 10^5^ microwells in an implanted surface of 25 mm^2^). To load neurons into these microwell scaffolds, we developed fixtures based on a PDMS stamp to create a seal around the coverslip. These fixtures contained a large reservoir to allow the implant to be loaded from a cell suspension pipetted on top of the scaffold, followed by capping and centrifugation under sterile conditions (Figure 1c).

**Figure 1:**
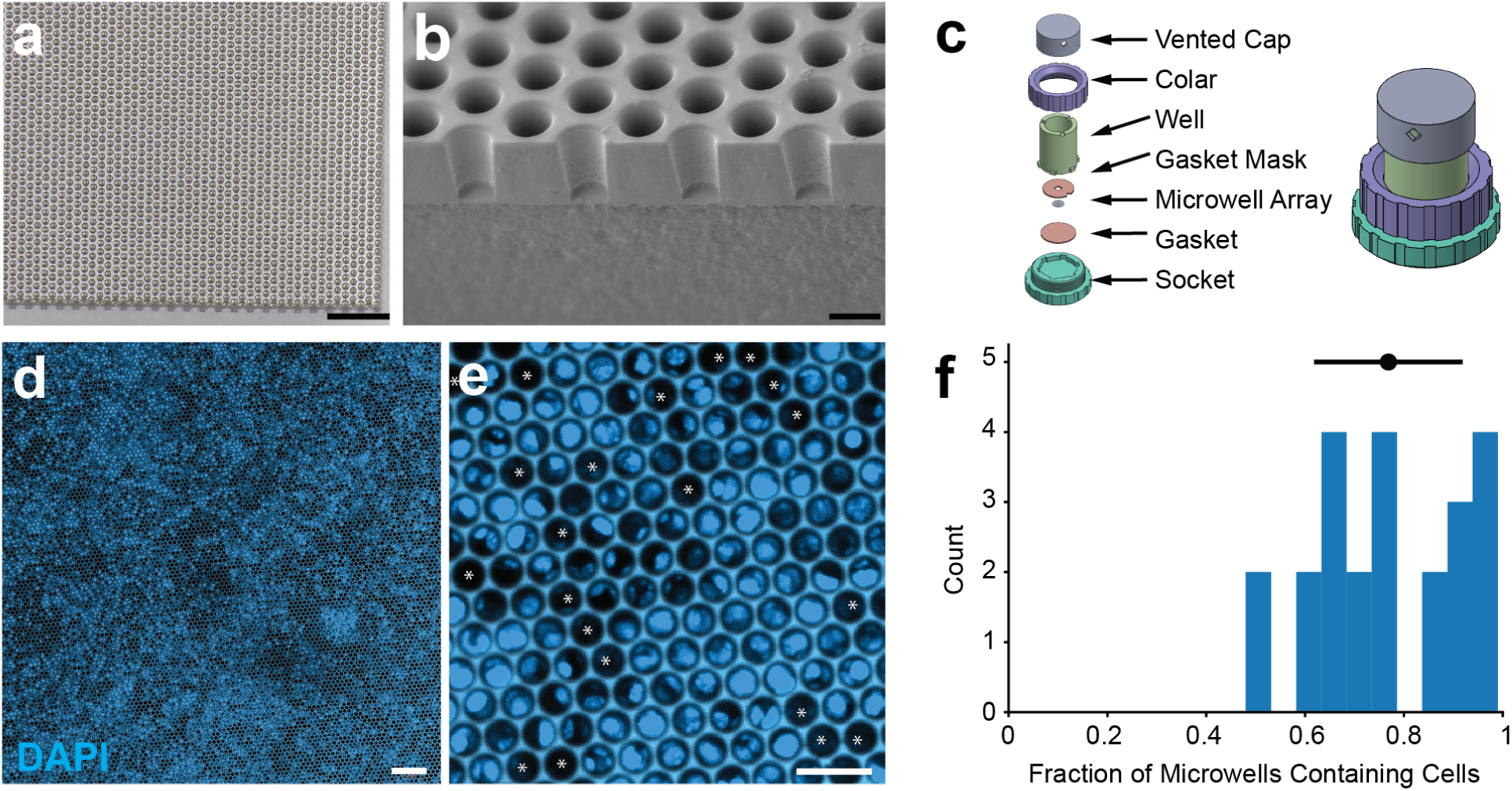
Fabrication and loading of microwell scaffolds. a. Light micrograph of a microwell scaffold bonded onto a glass coverslip. Scale bar is 100 µm. b. Electron micrograph of a cross-sectioned empty microwell scaffold taken at 45° tilt, sputtered with gold for imaging. Scale bar is 10 µm. c. Left, exploded view of the microwell loading fixture with labeled parts. Right, rendering of assembled loading fixture. d. Low zoom confocal micrograph of DAPI-stained loaded microwell scaffold taken 24 hours after loading. Scale bar is 100 µm. e. High zoom confocal micrograph of DAPI-stained loaded microwell scaffold taken 24 hours after loading. Scale bar is 25 µm. Asterixis indicates unloaded microwells. f. Histogram showing the fraction of microwells in individual scaffolds that were occupied by DAPI-positive staining 24 hours after loading (n=23 scaffolds, black circle and lines indicate mean and s.d.).

We performed embryonic dissections to obtain primary cortical neurons from C57/B6J mice at E14.5-E15.5^43^. This source of neurons had several advantages. Since these cells would be implanted back into other C57/B6J mice, this allowed for an autologous cell graft and obviated the need for any immunomodulatory therapies at this stage of development. Furthermore, at this embryonic stage, few inhibitory neurons have migrated into the cortex, so the transplanted cell population should be primarily glutamatergic excitatory neurons^44,45^. Indeed, we observed no teratomas or other overgrowth of the graft. To test our ability to load these embryonic primary neurons into the scaffolds, we performed dissections to obtain a cell suspension, and then used the fixtures to centrifuge cells into microwells. We allowed neurons loaded in microwells to recover in a CO_2_ incubator overnight, and then evaluated the array with light microscopy the next day. Imaging without counterstains could not reliably determine whether individual microwells contained cells (Supplemental Figure 1), so we fixed the arrays and used DAPI to determine how many microwells contained a cell nucleus. These experiments determined that on average we filled 77±15% (mean ± s.d., n=23 microwell scaffolds, Figure 1d-f). With this loading density, the mean implants contained ∼9 x 10^4^ neurons.

To implant the biohybrid device in mice, we performed a craniectomy followed by duratomy, placed the device cell side down on the brain, sealed the implant, and added a head post (Figure 2a, see methods). 2P imaging of implanted mice showed no visible sign of structural deterioration of the microwell scaffold even months after implantation. Microwell scaffolds exhibited slight autofluorescence, but were optically transparent and overall compatible with 2P imaging of the brain beneath the implants (Figure 2). We next transduced embryonic neurons with an adeno-associated virus (AAV) to allow visualization of the engrafted cells in microwell scaffolds using *in vivo* 2P microscopy. After primary cell dissociation and microwell scaffold loading, AAVs (CheRiff-eGFP (AAV2/2-Syn-CheRiff-eGFP) and/or jRGECO1a (AAV2/1-Syn-NES-jRGeco1a)^46,47^) were incubated with the cell loaded microwell scaffold overnight, followed by surgical implantation. Just prior to implantation, the biohybrid implant was washed in PBS to avoid introducing viral particles to the brain. Control experiments in which empty microwell scaffolds were incubated with AAVs in the absence of cells failed to observe any labeled cells in the brain, suggesting that this approach was sufficient to avoid virally transducing the brain (Supplemental Figure 2b).

**Figure 2:**
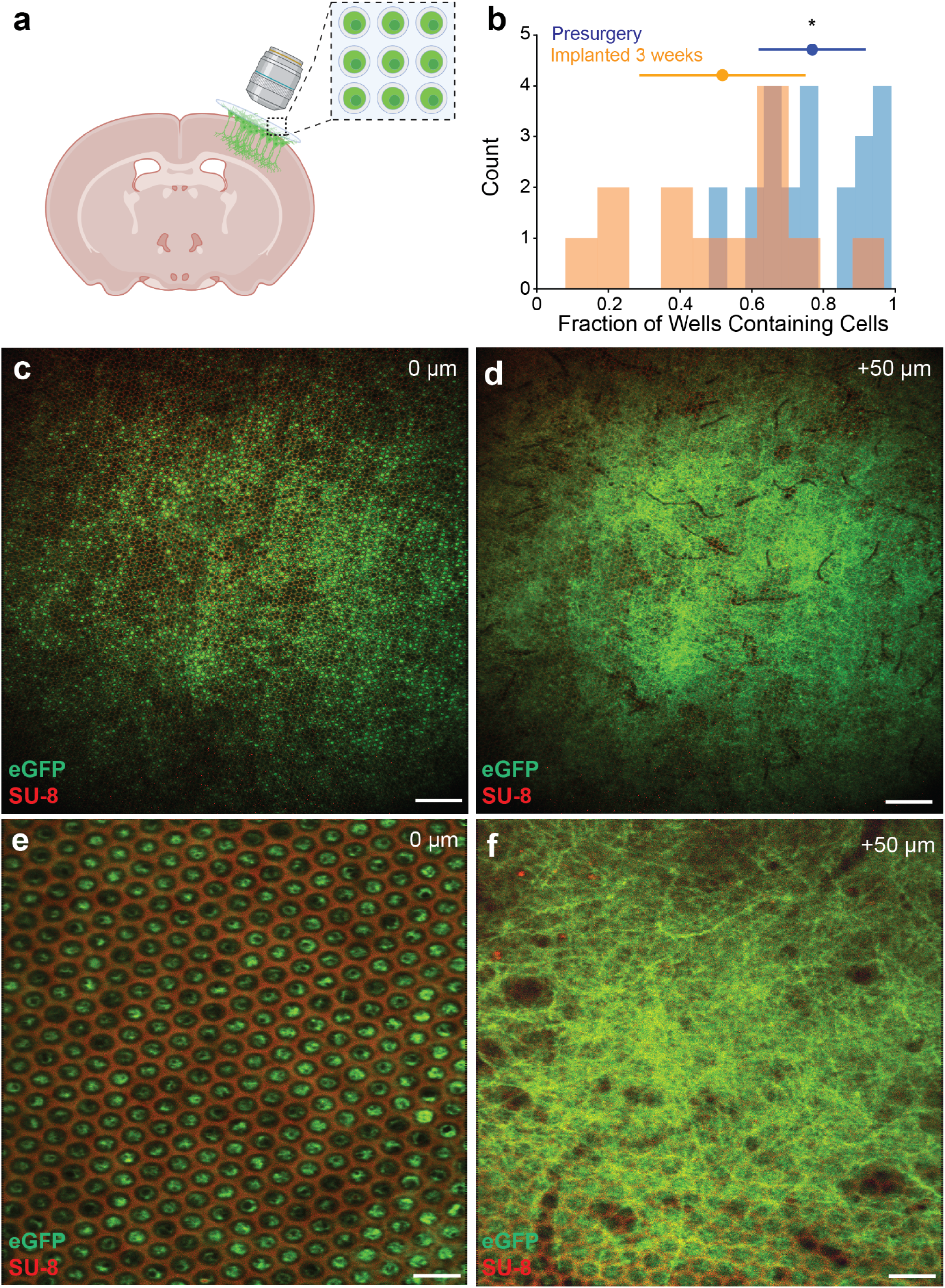
Robust survival and integration via cortical microwell scaffolds. a. Cartoon demonstrating experimental setup wherein mouse primary neurons are labeled with an AAV expressing CheRiff-eGFP and loaded into microwell scaffolds mounted on coverslips which are subsequently implanted onto the surface of the mouse brain. b. Histograms showing the fraction of microwells in individual scaffolds that were occupied by DAPI-positive staining 24 hours after loading (blue, n=25 scaffolds) and the fraction of microwells containing CheRiff-eGFP-labeled neurons via 2P microscopy approximately 21 days after surgery (orange, n = 13 implanted scaffolds). The distributions are significantly different (p = 0.0005, two sided t-test) c. Single 2P image from a representative mouse implanted with a microwell cell loaded scaffold approximately three weeks after surgery. The green channel shows virus-labeled CheRiff-eGFP positive engrafted cells, and the red channel collects autofluorescence from the implanted scaffold. The image plane is centered approximately on the center of the 20 µm cell scaffold. Scale bar is 100 µm. d. Single 2P image from the same mouse implanted in c, but here the image plane is centered approximately 50 µm below the cell scaffold above. Scale bar is 100 µm. e. As in c, 2P image centered on microwell plane, but at higher magnification and a different representative mouse three weeks after surgery. Scale bar is 25 µm. f. As in d, 2P image taken approximately 50 µm below the scaffold shown in e. Scale bar is 25 µm.

We next performed *in vivo* 2P imaging to assess graft survival and integration. Since cells did not express fluorescent proteins at the time they were surgically implanted, owing to the temporal delay in AAV expression after transduction^48^, we evaluated the success of the graft three weeks after surgery. At this time point, we found 52±23% (mean±s.d.) of microwells to be loaded with neurons expressing fluorescent protein (n=13 implants, Figure 2b-c, e). Although this constituted a significant reduction in loaded microwells relative to microwell scaffolds loaded for 24 hours before implantation (Figure 2b, p<0.005, two-sided t-test), it still represents robust survival of the graft, as central nervous system graft survival rates are typically less than 25%^18–20^. 2P imaging of tissue below the microwells made clear that engrafted neurons extended extensive processes into the superficial layers of cortex (Figure 2d,f, Supplemental Figure 2a). Imaging of microwells that had been incubated with virus and no cells or cells with no virus indicates that the fluorescence we observed in the superficial cortex was due to growth and integration of neurons expressing CheRiff-eGFP embedded in the implant (Supplementary Figure 2).

To assess whether grafted neurons were active, in a small subset of experiments (n=3 mice), we transduced neurons with a calcium indicator jRGECO1a prior to graft implantation. 2P imaging of these microwell scaffolds in awake mice head-fixed and running on a treadmill revealed spontaneous calcium events in these neurons three weeks after implantation (Figure 3a-b). Clustering of the pair-wise correlation coefficient between these calcium traces revealed multiple functional ensembles^49^, indicating that activity was not wholly correlated or uniform (Figure 3c). This data suggests that engrafted cells are able to mature and become active in the implant. We next assessed the degree of graft integration via histology. Unfortunately, dissecting the implant out of the skull without destroying the underlying cortex proved extremely difficult. Microwell scaffolds could not be left in place since they were adhered to a glass coverslip that prevented sectioning. When they were removed, they routinely fractured and pulled large chunks of cortical tissue out with them. Examination of explanted arrays indicated that cells remained in microwells even weeks after implantation. The presence of blood vessels in the tissue pulled from the brain during explant revealed tight mechanical coupling between the cortex and implanted cells in the microwell scaffold (Figure 3d). Although the damage to cortex caused by explant made systematic histology impossible, in some animals we were able to obtain some sections near the implant site and were able to confirm our observations via 2P microscopy of robust integration in the superficial cortex. Additionally, we were able to observe axons throughout the layers of the cerebral cortex, suggesting that neurons implanted above layer 1 are able to extend axons deep into the brain (Figure 3e).

**Figure 3:**
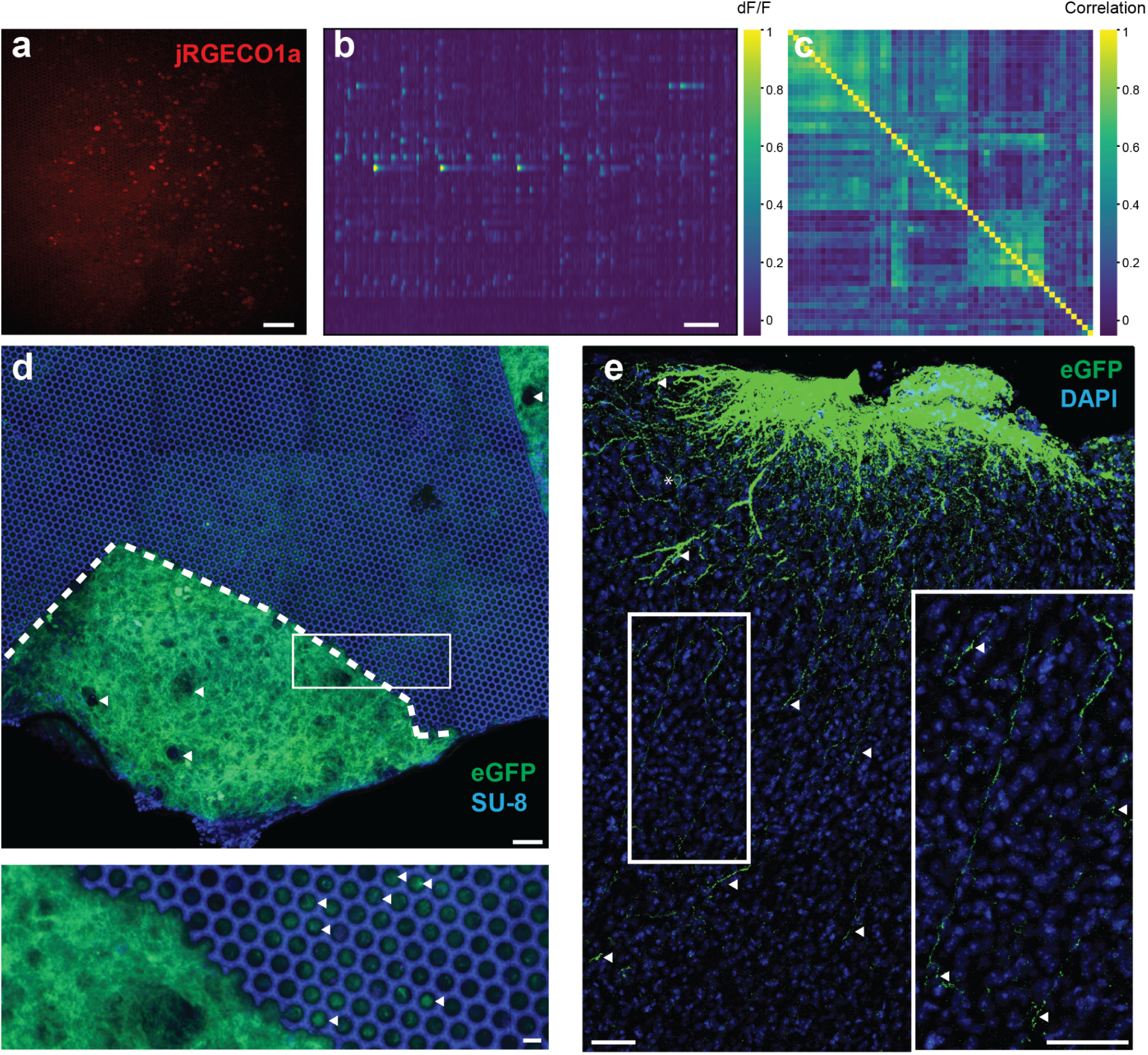
Cells in microwell scaffolds mature and integrate. a. Average image created from 2P functional imaging of implanted cells in microwell scaffold expressing jRGECO1a. Scale bar is 25 µm. b. Representative df/f traces from 30Hz 2P imaging of jRGECO1a-expressing cells loaded in a microwell scaffold. Figure shows spontaneous activity from a head fixed mouse free to run on a wheel. Scale bar is 10 seconds. Colorbar shows df/f values from 0 to 1. c. Representative dendrogram showing pairwise correlation coefficients from spontaneous activity of cells implanted in microwell scaffolds. Colorbar shows correlation coefficients from 0 to 1. d. Confocal micrograph (single plane) of representative explanted microwell scaffolds months after implantation. Cells in this experiment were transduced by AAV expressing CheRiff-eGFP (green) and the scaffold’s intrinsic fluorescence dominates the blue channels. Explanting microwell scaffolds was usually very destructive; in this example a region of the microwell scaffold outlined in the white dashed line fractured, but the tissue beneath was sufficiently adhered to the scaffold that it pulled up from the brain. The presence of blood vessels (white arrows) indicates that the brain was tightly adhered to the graft. The scale bar is 50 µm. The inset below shows loaded cells still in their microwells (white arrows), scale bar 10 µm. e. Confocal micrograph (maximal intensity projection) of representative region of cortex beneath an explanted microwell scaffold removed months after implantation. Cells in this experiment were transduced by AAV expressing CheRiff-eGFP (green), nuclei are labeled with DAPI (blue). Dense processes are observed in the superficial cortex, but putative axons are visible deep in the cortex (white arrows; inset). Scale bars are 50 µm and 25 µm in the inset. Asterix indicates the cell body in the cortex.

Since our engrafted neurons survive and exhibit spontaneous activity, we next sought to determine if the graft was functionally integrated with the host brain. To accomplish this, we elected to design a behavioral task in which the mouse would be required to use the activity of the engrafted neurons to obtain rewards. In this task, an animal initiates a trial by activating a center nose-poke port, and then must report whether optogenetic stimulation occurred by activating a left or right port to obtain a water reward^39,40^ (Figure 4a-b, see methods). To validate the task, we first trained two control cohorts. Positive control animals were injected with an AAV encoding an excitatory opsin (hChR2-eGFP (AAV1-eSYN-hChR2(H134R)-eGFP) (n=5) or CheRiff-eGFP (n=2)) in primary somatosensory cortex (S1), while negative control animals were injected with an AAV expressing a calcium indicator (jRGECO1a (n=4)), which should not alter neural activity in response to light. Four weeks later, we implanted a cranial window, screened for expression, and installed a fiber optic ferrule attached to an LED. We observed that 86% (6 of 7) positive control mice injected with the excitatory opsin reached criterion performance (d’ > 1.25) within the three week training window (9.5±3.5 days to criterion mean±s.d.). In contrast, none of the mice in the negative control cohort injected with calcium indicator reached criterion during the three weeks of training. This confirms that optogenetic activation of neurons using a fiber-coupled LED was required to successfully complete the task, and that animals were unable to use the presence of light, sound, heat, or other external stimuli as cues to complete the task (Figure 4c-e).

**Figure 4:**
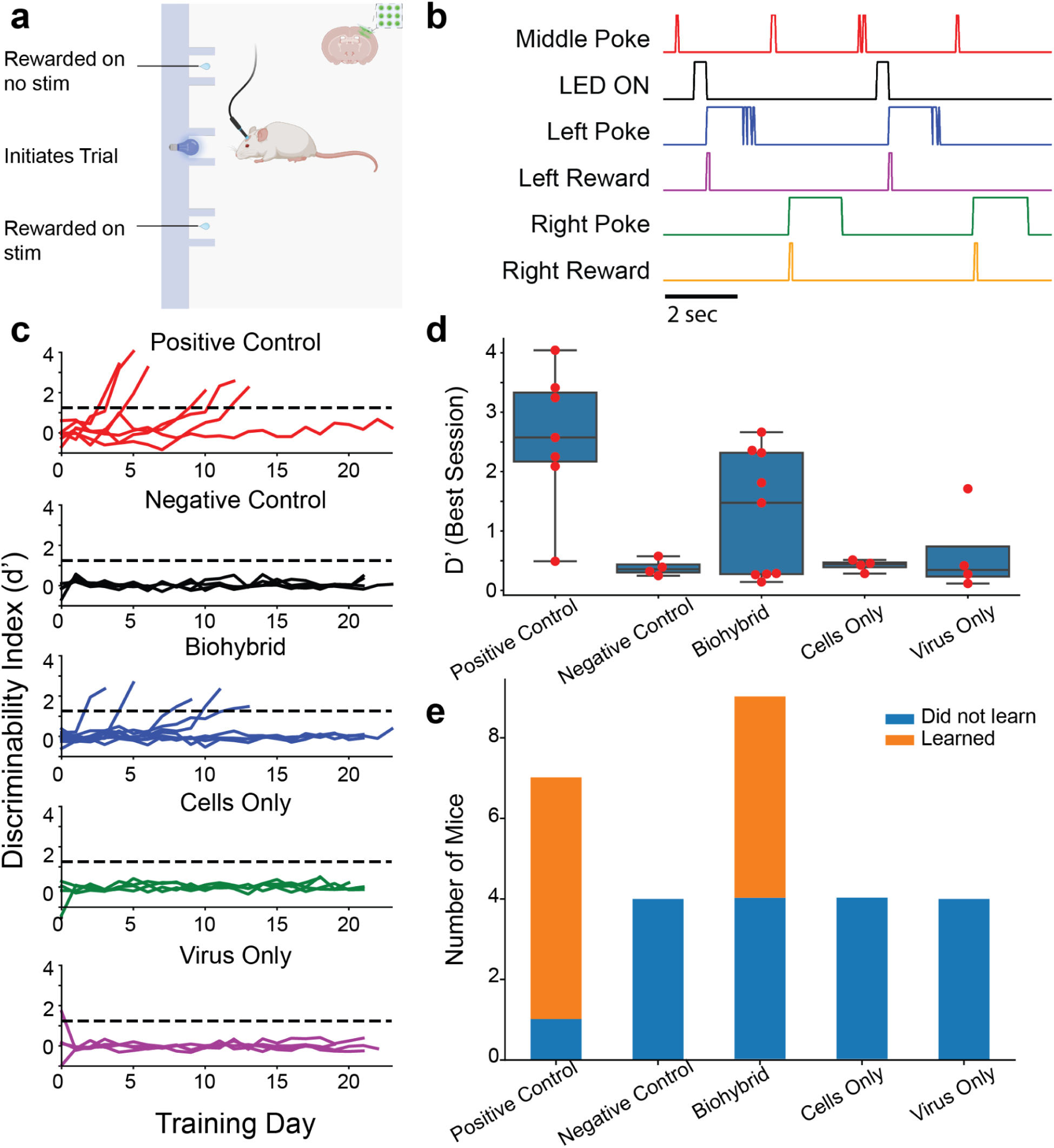
Optical stimulation of the biohybrid implant guides goal directed behavior. a. Cartoon illustrating task in which mice implanted with biohybrid implants learn to detect stimulation of the grafted cells to gain rewards. Briefly, trials are initiated by breaking an IR beam at the center nose poke, which triggers masking lights on every trial. Rewards are administered after breaking the beam on the right port on trials where there is no stimulation and on the left port on trials in which there is optogenetic stimulation of the implant. b. Representative sequence of four trials, all correct, in which the mouse alternates between the right and left ports on stimulation and catch trials to obtain water rewards. c. Plots showing training day (*x-*axis) versus discriminability index d’ on the *y*-axis. On each plot a dashed black line indicates criterion performance. Each colored line is the score from an individual animal on a given training day. Subplot are positive control (AAV-excitatory opsin injected into cortex, n=7, red), negative control (jRGECO1a AAV injected into cortex, n=4, black), biohybrid implants (engrafted neurons on microwell scaffolds with CheRiff-eGFP, n=9, blue), cells only (microwell scaffolds loaded with cells and no AAV, n=4, green), or virus only (microwell scaffolds incubated with CheRiff-eGFP, n=4, magenta). Note that in each case animals were trained for 21 sessions or until they achieved criterion performance on two consecutive days. d. Box and swarm plots showing the discriminability index for the best session for each animal in each condition. Each dot corresponds to a single animal, and the boxplots show mean and interquartile range (p < 0.005, one-way ANOVA). e. Stackplot showing the total number of animals in each group that passed criterion performance (orange) or failed to pass criterion performance (blue) during the 21-session training window (p<0.005, Chi-Squared test).

We then trained a cohort of animals implanted with our biohybrid implant to complete the task (n=9). These implants were loaded with neurons incubated overnight with the excitatory opsin CheRiff-eGFP. Three to four weeks later, animals were screened for expression, implanted with a fiber optic ferrule and then entered behavior training. Five of nine mice in this cohort passed criterion performance in the three weeks training period (Figure 4c-e, not statistically significant relative to the positive control group, Fisher Exact Test p=0.3). For animals that did reach criteria, time to criterion performance was identical to the positive control group (9.6±4.0 days to criterion mean±s.d., Supplementary Figure 3, p=0.95, t-test). To control for the possibility that residual AAV in the microwells transduced neurons in the brain, we trained an additional control group where we implanted microwell scaffolds that had been incubated with the excitatory opsin CheRiff-eGFP, but not loaded with cells. We did not observe fluorescence in these mice via 2P microscopy (Supplementary Figure 2b), and none (0 of 4 mice) of the animals in this group achieved criterion performance (Figure 4c-e). This suggests that if any residual AAV did transduce cells in the host brain, it was not sufficient to drive detection of the light stimulus. Additionally, no (0 of 4) mice that received biohybrid implants with no AAV incubation achieved criterion performance, indicating that neuron engraftment alone is not sufficient to allow mice to detect optical stimuli (Figure 4c-e). Since we imposed a trial structure that allowed a variable response time after stimulus, we were able to retroactively assign a bit-rate to each mouse during active trial time. Remarkably, biohybrid mice achieved a positive bit rate similar to positive controls and sustain bit rates as high at 0.7 bits / seconds (Supplementary Figure 3, best session performance for biohybrid rate averaged 0.25±0.24 bits per second mean±s.d., n = 9). Overall, this data supports the hypothesis animals implanted with biohybrid devices above cortical layer 1 have access to information originating from the activity of the graft, and are able to use that information to guide goal-directed behavior.

## Discussion

This study provides a proof-of-concept demonstration that an optogenetic biohybrid neural implant positioned on the cortical surface can be activated by light and used for goal directed behavior. The implant consists of tens of thousands of neurons positioned on the top of cortical layer 1 in microwell scaffolds. Since the number of surviving engrafted cells is roughly equivalent to the number of neurons in a cubic millimeter of mouse cortex^50^, it is perhaps not surprising that the animal is able to detect stimulation of the graft, as the stimulus could potentially generate hundreds of thousands of action potentials. Indeed, detection of optogenetic stimulation of the brain has been reported to require as few as several hundred neurons^39^, so our experiments do not provide a clear readout of the degree of graft integration with the cortex. While it is likely that the graft forms chemical synapses with the host brain that are used to read out the state of the engrafted cells, we cannot rule out the possibility that the mouse uses other activity-dependent cellular mechanisms to complete the task. Regardless, since neither implantation of the empty virus-washed scaffold nor opsin-negative cells is sufficient to complete the task, the animals clearly have some way to use the graft to detect activation of the LED. Mice can learn to act on optical stimulation of many brain areas^40^, and similarly to optogenetic stimulation of various brain areas, optogenetic activation of engrafted cells likely provides a perturbation to on-going activity that the animal is able to monitor. Not all of the engrafted mice learned to perform the task. Opsin expression and survival of the graft cannot explain this result, as animals were pre-screened for expression before entering training, and the number of observed opsin-expressing cells was not correlated with task performance (Supplementary Figure 4a). However, the engrafted animals that did learn the task performed as well as the positive control group and also learned as quickly (Supplementary Figure 4b).

Surprisingly, our grafts exhibited very high rates of neuronal survival (∼50%). Engraftment of neurons via microinjection results in only a small fraction of surviving cells across a wide range of neuronal cell types and brain regions^18–21,51,52^. Apoptosis after engraftment^53^ is thought to be driven by hypoxia and immune insult associated with disruption of the blood brain barrier^21^, and occurs in the first few days after engraftment^18,52^. By placing our implant on the cortical surface, we minimize the disruption of the vascular network, and therefore avoid the major mechanism of cell death after transplantation. Additionally since our graft is one cell-layer thick and spread evenly across the cortex, we avoid the deleterious microenvironment thought to be present in the center of a bolus of injected cells. Since quantification of microwell loading required fixing cells, we were unable to measure the number of microwells loaded with neurons pre- and post-implantation to quantify the number that survived transplantation. Similarly, use of a viral transgene to visualize our graft neurons meant that we were not able to measure graft survival in the first several weeks after surgery. Some apparently empty microwells likely have healthy neurons that were not transduced by the virus, or may have been occupied by cells unable to express proteins downstream of the synapsin promoter. We note that the use of allogeneic embryonic primary neurons in this study may be a key factor governing graft survival, and is not an option for translational approaches.

Although placing the biohybrid device on the cortical surface provided key advantages for graft survival and future integration with devices, it caused unique challenges for graft integration. Specifically, in order to integrate with the brain, neurons needed to extend processes through the pia mater, which specifically functions to prevent infiltration into the brain^54^. Indeed, most grafted cell processes were located in the superficial cortex, although sparse axons were detected throughout the cortex (Figure 3e). Occasionally, labeled cell bodies were detectable in the superficial cortical layers. We cannot tell if these are engrafted neurons that migrated out of their microwells or if they are the result of residual AAV sparsely transducing the cortex. Our behavioral control cohorts suggest that even if these labeled cells are the result of residual virus, they are not sufficiently numerous to allow the animal to perform the detection task. The presence of numerous neurons in microwells months after engraftment suggests that migration is rare, but we cannot categorically rule out some degree of migration out of the microwells and into the cortex.

The ability to learn to detect graft stimulation implies that an animal has access to the information conveyed by the engrafted neurons’ activation and can use it to guide a goal directed behavior, in this case to obtain water rewards. Thus, this report demonstrates the first example of a biohybrid implant’s use in a brain computer interface task. In this report, we elected to leave the top of the implant optically transparent so as to allow live imaging of neurons in the graft, and behavior was driven by commercial fiber-coupled LEDs. However, we and others have fabricated high density µLED displays^55^ at similar pitch to the microwell scaffolds. Future versions of a biohybrid implant could allow pixels to be aligned to microwells to allow stimulation at near single-cell resolution. Although numerous challenges exist towards translating biohybrid neural interfaces for extremely high bandwidth BCIs, this study represents a proof-of-concept demonstration that such devices can in principle be used to guide goal directed behavior.

## Methods

### Animals

C57/B6J male and female mice obtained from Charles River were used in this study. Mice were maintained on a 12 hour light-dark cycle. All animal procedures were carried out with the approval of Science Corporation’s institutional animal care and use committee.

### Fabrication of microwell scaffold implants

Microwell scaffolds were fabricated on 100 mm fused silica wafers using photolithography. In preparation for lithography, the wafers were cleaned using a spin rinse dryer, dehydrated for 15 minutes on a 200℃ hotplate, and exposed to O_2_ plasma in a Technics RIE for 2 minutes at 50W and 300mTorr. To improve microwell adhesion, a thin layer of SU-8 2001 (Kayaku Advanced Materials) was spun at 3000 rpm and UV flood exposed to anchor the subsequent thicker layer. To achieve a microwell height of 15 µm, SU-8 2010 photoresist (Kayaku) was spun at 3000 rpm for 30 seconds, soft-baked at 95℃ for 3.5 minutes on a hotplate, and exposed with a Karl Suss MAB6 mask aligner using contact lithography and a total UV dose of 294mJ/cm^2^. Exposure was done utilizing a long pass filter to filter wavelengths below 360 nm. We completed a post exposure bake of 4 minutes at 95 ℃ and developed for 3.5 minutes in SU-8 Developer (Kayaku). To improve biocompatibility^56^, a UV flood-exposure was used with a total UV energy of ∼1.86 J. Finally, the wafers were hard baked by holding at 95℃ for 5 minutes and then ramping to 200℃ for 2 minutes on a hotplate. Wafers were then coated in SPR220-7 photoresist as protection for dicing and were diced into 5×5mm die using a Disco DAD3240. The dies were then demounted from tape and rinsed in acetone followed by isopropyl alcohol to remove the protective photoresist.

Individual microwell die were then mounted to 7mm diameter glass coverslips using a small drop of Epotek 301 epoxy and cured in an oven at 65℃ for 2 hours.

Loading fixtures were fabricated using 3D printing and CO_2_ laser cutting techniques. All components not in contact with the biological solution were FFF (Fused Filament Fabrication) printed (Raise3d ProPlus printer, commercial PLA+ filament [ANYCUBIC PLA+ Gray or equivalent]) . Nominal hot-end temperature was 205℃, and the bed was heated to 50℃. The bed interface was treated with a film of PVA (Elmer’s All Purpose Glue Stick) to aid in part-to-bed adhesion. After printing, all FFF parts were scrubbed and soaked in deionized water for a minimum of one hour. Gaskets were made from FDA 177.2600 approved silicone sheets. A CO_2_ laser cutter (Full Spectrum Muse) was used to cut the gasket shape. After laser cutting, all gaskets were soaked in deionized water for a minimum of one hour. The well was fabricated using a photopolymerization printer (FormLabs 3B, BioMed Clear resin) according to manufacturer’s recommendations, followed by a one hour soak in deionized water.

### Preparation of loaded biohybrid scaffold

Microwell scaffolds were sterilized in 70% ethanol and left to dry under UV light exposure overnight in a biosafety cabinet. Prior to loading neurons, microwell scaffolds were loaded with 0.1 mg/mL poly-d-Lysine (Gibco) and 0.1 mg/mL Laminin (Millipore Sigma, L2020) in DI water for at least one hour. Following incubation, arrays were washed three times with sterile water and placed in a CO_2_ incubator in a sterile petri dish.

Embryonic cortical neurons were prepared essentially as described^43^. Briefly, pregnant C57/B6 mice at embryonic day E14.5-E16.5 were sacrificed via CO_2_ euthanasia, sprayed down with 70% ethanol then transferred to a sterile dissection hood. The uterine sac was transferred to ice cold dissection media (HBSS supplemented with 100 mM MgCl_2_*6 H_2_O, 100 mM HEPES, and 15 mM Kyneuric Acid adjusted to pH 7.2). Embryos were removed from the uterine sack, and the brains were dissected out under a stereoscope using a pair of fine forceps. The meninges were removed, and the cortices collected in ice cold dissection media. After all cortices were collected, the tissue was incubated in papain solution (dissection media supplemented with 2.5 mM L-cysteine and 100 units of Papain (Millipore Sigma) in a CO_2_ incubator for 5-7 minutes. The tissue was carefully washed three times with prewarmed light inhibitor solution (dissection media with 6 mg/mL bovine serum albumin and 6 mg/mL Trypsin inhibitor followed by three washes with heavy inhibitor solution (dissection media with 60 mg/mL bovine serum albumin and 60 mg/mL Trypsin inhibitor). Following the heavy inhibitor wash, the tissue was washed once with prewarmed Neurobasal medium and then triturated into single cell suspension in Neurobasal+ media (neurobasal supplemented with 1% Penicillin-Streptomycin, 1% Glutamax, and 2% B27 supplement. Cell concentration was counted and adjusted to 1e6 neurons / mL and kept on ice.

Microwell scaffolds were removed from the incubator, and assembled in a loading fixture in the biosafety cabinet using sterile technique. The neuron suspension was transferred into a loading fixture, covered, and centrifuged at 300 RCF for 5 minutes at room temperature. The loaded microwell scaffold was incubated in a CO_2_ incubator for two hours, and then removed from the fixture in a biosafety cabinet two hours later and placed into a standard cell culture dish with fresh prewarmed neurobasal+. In some experiments, AAVs (CheRiff-eGFP: AAV2/2-Syn-CheRiff-eGFP (Biohippo #BHV12400513, titer: 2.37e12 vg/mL), jRGECO1a: AAV2/1-Syn-NES-jRGeco1a (Biohippo #BHV12401534, titer: 5.85e12 vg/mL)) were added to the media for an overnight incubation prior to surgical implantation the next day.

### Viral Injections

For all surgical procedures, mice were anesthetized with isoflurane (2%), and administered 2 mg/kg of dexamethasone as an anti-inflammatory and 0.05 mg/kg Ethiqa as an analgesic. Animals were maintained on isoflurane on a heating pad and head fixed in a stereotactic apparatus (Kopf). After sterilizing the incision site, the skin was opened, and a small burr hole was drilled over S1 (ML 2mm, AP +2mm from Lambda) using a 0.24 mm drill bit (Busch). 750–1000 nL of virus was injected using a micro syringe pump (Micro4) and a Wiretrol II glass pipette (Drummond) at a rate of 25 nL/second 3-4 depths 1500-500 μm below the brain surface. After the injection was complete, the needle was held in place for several minutes before suturing the scalp closed. Viruses used: AAV2/2-Syn-CheRiff-eGFP (Biohippo #BHV12400513, titer: 2.37e12 vg/mL), AAV1-eSYN-hChR2(H134R)-eGFP (Biohippo #BHV12400006, titer: 1.6e13 vg/mL), AAV2/1-Syn-NES-jRGeco1a (Biohippo #BHV12401534, titer: 5.85e12 vg/mL). days after virus injection, mice were anesthetized, and a cranial window and headplate was installed over the injection site. The scalp was removed, and the fascia retracted. Following application of Vetbond (3M) to the skull surface, a custom stainless steel headplate was fixed to the skull with two dental cements: Metabond (C&B) followed by UV-cure acrylic (Flow-IT ALC). After the dental cement dried, a 3 mm diameter craniotomy over the left S1 was drilled, and residual bleeding stopped with repeated wet-dry cycles using sterile hypotonic saline, gauze, and Gelfoam (Pfizer). A window plug consisting of two 3mm diameter coverslips adheredto the bottom of a single 5 mm diameter coverslip (using Norland Optical Adhesive #71) was placed over the craniotomy and sealed permanently using Metabond and UV-cure acrylic. Animals were allowed to recover in a heated recovery cage before being returned to their home cage. Approximately three days after surgery, animals were habituated to head fixation under a freely moving circular treadmill and screened for virus expression under an upright epifluorescence microscope (Thorlabs Cerna, 10x mitutoyo objective 0.28 NA).

### Microwell Scaffold Implantation

Mice were anesthetized with isoflurane (2%), and administered 2 mg/kg of dexamethasone as an anti-inflammatory and 0.05 mg/kg Ethiqa as an analgesic. The scalp was removed, the fascia retracted, and the skull lightly etched. A 5 mm diameter craniotomy was drilled over S1, and residual bleeding stopped with gelfoam (Pfizer). A duratomy was performed using a von graefe knife, and the dura carefully retracted using fine forceps. Residual bleeding was again stopped by gelfoam. Microwell scaffolds loaded with cells the day before surgery were picked up with forceps, and gently washed three times with sterile neurobasal media or PBS before being inverted and placed into the craniotomy. Downward pressure was applied using a stereotaxic attachment, and the interface between the 7 mm outer coverslip and the skull filled with Metabond. After the metabond dried, a custom stainless steel headplate was fixed to the skull using Metabond and UV-cure acrylic. Animals were allowed to recover in a heated recovery cage before being returned to their home cage. Animals were evaluated using a 2P microscope approximately three weeks after surgery.

### Ferrule Implantation

Following screening for expression and/or engraftment, mice were anesthetized with isoflurane (2%) and mounted in a stereotaxic frame. A 400 µm fiber optic cannula (Thorlabs CFMLC14L02, 0.39 NA) was secured to the top of the glass cranial window using several drops of UV-cured optical acrylic (Norland 71). The implant was then covered in UV-cure acrylic, cured, and then covered with metabond mixed with black oxide to make it optically transparent. Animals were allowed to recover from anesthesia in a heated recovery cage before being returned to their home cage, and they began water restriction for behavior several days later.

### 2P Imaging

For 2P imaging, mice were head-fixed on a freely spinning running wheel under a Nixon 16x-magnification water immersion objective (N16XLWD-PF) and imaged with a custom built 2P resonant scanning microscope within a darkened box. The running wheel setup was mounted on a Hexapod (Physik Instrumente) which allowed the plane of the implant to be roughly matched to the microscope imaging plane. Structural imaging was performed using ScanImage5 (Vidrio), and z-stacks were acquired by moving the objective using a piezo actuator (PFM450E, Thorlabs). Imaging was performed at an average power of 100 mW at 930 nm using a Coherent Discovery femtosecond laser. For functional jRGECO1a imaging, single plane imaging was performed at 30 Hz using a wavelength of 1040 nm at an average power of ∼50 mW. Images were analyzed without motion correction, since the imaging plane was stable and the hexagonal lattice structure of the microwells, which were always visible due to faint fluorescence, disrupted these operations and made the resulting motion corrected stacks far worse than the uncorrected stacks. Calcium sources were identified morphologically using CellPose^57^ on an average image. Average calcium intensity values were converted to dF/F by baselining using the 10th percentile of intensity values of each calcium source. No neuropil subtraction was performed since there was no neuropil in the imaging plane; the space between somas was occupied by the microwells, which were visually confirmed to not contaminate calcium source.

### Histology

Animals were deeply anesthetized using Ketamine and transcardially perfused with cold PBS followed by 4% paraformaldehyde. After perfusion, the head was removed and we attempted to remove the metabond holding the implants in place using a dental drill. Implants were removed using forceps, but we often observed strong adhesion between the implant and the brain. This adhesion resulted in numerous tissue rips and tears on the implant’s surface, including numerous instances in which large chunks of brain under the cranial window were pulled out when the window was removed. After removing the implant, the rest of the brain was dissected out of the skull, and then post-fixed for 24 hours, followed by cryoprotection in sucrose gradients. After the brain sank in 30% sucrose it was snap frozen in OCT using a isopentane liquid nitrogen bath. Samples were left in the -20C freezer overnight. Using a Leica cryostat, 50 µm cryosection retinal slices were made and mounted onto Fisherbrand™ Superfrost™ Plus Microscope Slides (Fisher Scientific Cat No. 12-550-15). Samples on slides were then blocked in a solution containing 10% donkey serum (DS), 0.1M Glycine, 0.3% Triton X-100 in PBS for 1 hour. Primary antibodies were diluted in 1% DS in PBS at room temperature overnight. Samples were washed in PBS three times for 5 minutes, then incubated for 2 hours at room temperature with secondary antibodies diluted in PBS. Samples were washed in PBS three times for 5 minutes, then 4’,6-Diamidino-2-Phenylindole, Dihydrochloride (DAPI) (D1306 Thermofisher Scientific) (1:1000) was added to all the samples to stain cell nuclei and incubated at room temperature for 5 minutes. All samples were washed again with PBS for 5 minutes. Fluorescent images were captured using a ZEISS LSM 980 confocal microscope. Micrographs were captured using the Plan-Apochromat 10x/0.45 air objective (Carl Zeiss Microscopy, Jena, Germany). ZEN Blue 3.5 software was used to perform stitching of tilescans and maximum intensity projections of Z-stacks.

### Optical Stimulation Task

After surgery, mice were placed on a high fat diet for at least one week. After baseline weight was established by measuring daily for three consecutive days, animals were restricted to 1 mL of water daily for 5-9 days before beginning behavior training. Throughout training, mice were weighed daily and given supplemental water if their weight fell below baseline for 1 day, and were given access to free water if their weight didn’t recover to at least 80% of baseline weight after supplemental water.

The optical microstimulation task was conducted in a 9 x 9 inch box in which the animal was allowed to freely move. On one side of the box, three nose poke ports were installed. Each port contains an IR beam break. The center point contains masking light LEDs and the left and right ports contain a lick spout controlled by two solenoid valves (Nresearch) connected to a water reservoir calibrated to set each reward at 50 uL of water. Experimental control was conducted with an microcontroller (Arduino Due) mated to a custom circuit board. Data was streamed via serial port to a PC that allowed the experimenter to control the task via custom software. Prior to a session, the cover on animals’ implanted optical fiber ferrule was removed, and a 400 µm optical fiber was inserted into the ferrule. The fiber was commutated by a rotary joint (Thorlabs RJPSF2) and attached to a Thorlabs LED (470 nm) controlled by a Thorlabs LEDD1B driver. Optical power throughput was measured using a power meter before surgical implantation, and behavior was run at levels that corresponded to 5 mW of optical power out of the ferrule before surgical implantation. In some high performing animals, we reduced the power to 1 mW and they were still able to perform the task, but we did not systematically test the effect of optical power on performance.

Mice were introduced to the task in three phases. In phase 1, the animals learned to navigate the box with an optical fiber attached to the head and to collect water rewards from both the left and right port. In this phase, triggering the center nose port had no effect, but triggering the left or right nose ports resulted in an immediate water reward, with a variable 1-5 second cooldown period before the next reward was available. Animals are advanced to phase 2 as soon as they obtain 15 rewards from both ports in 10 minutes, demonstrating that they have the ability to trigger the IR beam break and obtain water.

In phase 2, animals learned to initiate a trial in the center port and collect reward on the left or right ports. In this phase, rewards were only available on the left and right ports after the center beam had been broken. Since the mouse was now able to associate the noise from the solenoid valve with the availability of reward, each center poke triggered an immediate reward available on either the left or right port (changed randomly each trial). If the animal poked the correct port within 10 seconds, additional reward was given from that port. After 20 successful trials on each side in a session, animals were advanced to phase 3.

Phase 3 is the complete task. Mice initiated trials by poking the center port, which triggered the immediate activation of masking lights (also activated in earlier phases) as well as optogenetic stimulation on some trials (10 pulses at 10 ms, 20 Hz). Reward was available on the left side for optogenetic stimulation or the right side for no stimulation. To avoid rewarding a strategy where a mouse could simply go to one side for consistent reward on 50% of trials, trial type was pseudorandomized to adjust for bias every 30 trials to encourage mice to engage with both the left and right ports. Trials were scored as HIT trials if it successfully reported optogenetic stimulation, MISS if it failed to select the correct report after stimulation, FALSE ALARM (FA) if it selected the stimulation port after a catch (no stimulation) trial, or Correct Reject (CR) if it selected the right port during a catch trial.

For each behavioral session, which lasted 30 minutes or ∼100 trials, we calculated the discriminability index d’. We compensated for trial types that didn’t occur by adjusting zeros by resetting trial types with zero events to 1 - 1/2N, where N = number of trials. We then calculated hit rate (HR; hits / hits + misses) and false alarm rate (FAR; FA / FA + CR). We z-scored these using the inverse cumulative distribution function (CDF) of the standard normal distribution, and subtracting to find d′=Z(HR)−Z(FAR). Criterion performance was achieving d’ > 1.25 for two consecutive sessions, after which training was stopped. All training was halted after 21 sessions, or 8 sessions longer than the slowest-learning animal in the positive control group, and these animals were deemed to have not learned the task. Training occurred 5 or 6 days a week, with a 1 mL of free water administered on break days. Bit rate was calculated on a per-session basis using Wolpaw’s definition^58^ for N=2 choices, bit_rate = (p + (1 - p) * np.log2(1 - p)) / T where p is average success rate and average trial duration.

## Supplementary Figures

**Supplementary Figure 1:**
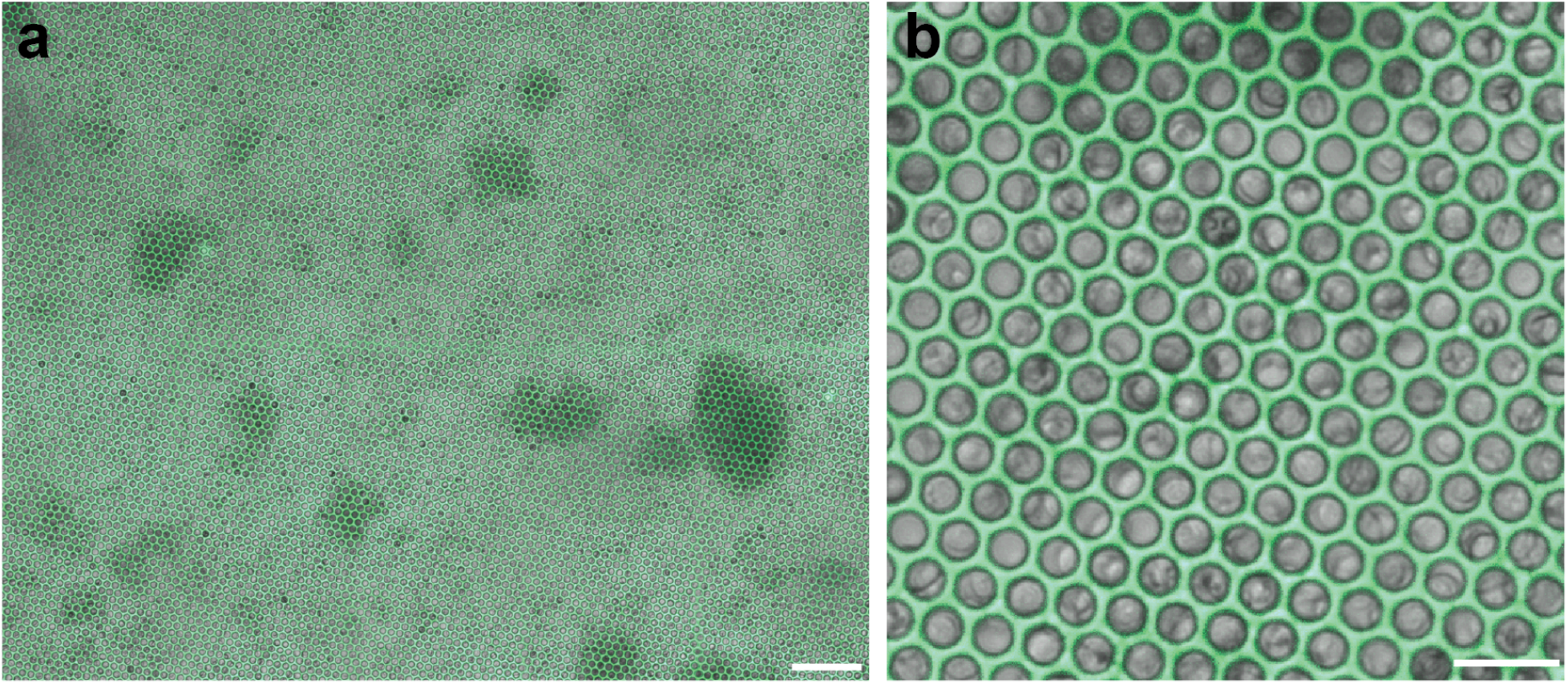
Live imaging of loaded microwell scaffolds pre-implantation. a. Confocal micrograph showing mouse embryonic neurons (gray, transmitted PMT) in SU-8 microwell scaffolds (green, scale bar 100 µm). b. Higher zoom confocal micrograph showing mouse embryonic neurons (gray, transmitted PMT) in SU-8 microwell scaffolds (green, scale bar 25 µm).

**Supplementary Figure 2:**
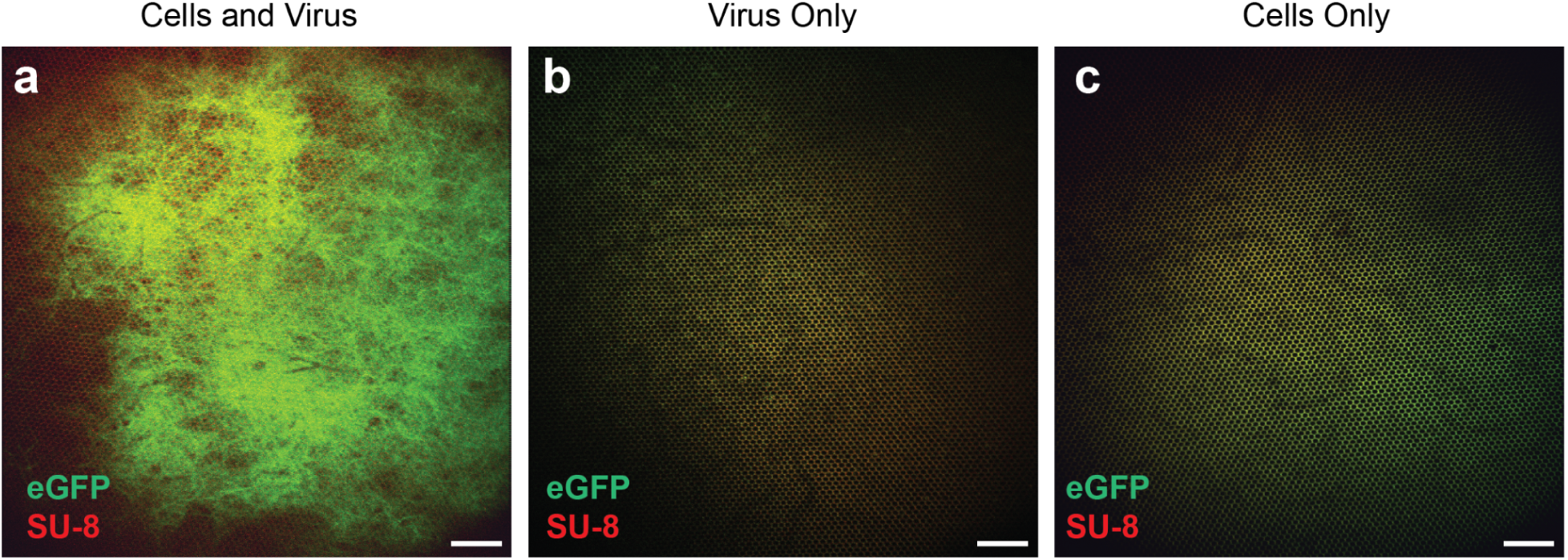
2P Imaging of control microwell conditions. a. Maximal intensity projection of a 2P image stack from a representative mouse implanted with a microwell scaffold (red) containing neurons incubated with an CheRiff-eGFP virus for 24 hours before implantation and then washed pre-surgery. Image is taken three weeks after surgery. Scale bar is 100 µm. b. Maximal intensity projection of a 2P image stack from a representative mouse implanted with a microwell scaffold (red) without cells incubated with an CheRiff-eGFP virus for 24 hours before implantation and then washed pre-surgery. Image is taken three weeks after surgery. Scale bar is 100 µm. c. Maximal intensity projection of a 2P image stack from a representative mouse implanted with a microwell scaffold (red) loaded with cells without any virus. Image is taken three weeks after surgery. Scale bar is 100 µm.

**Supplementary Figure 3:**
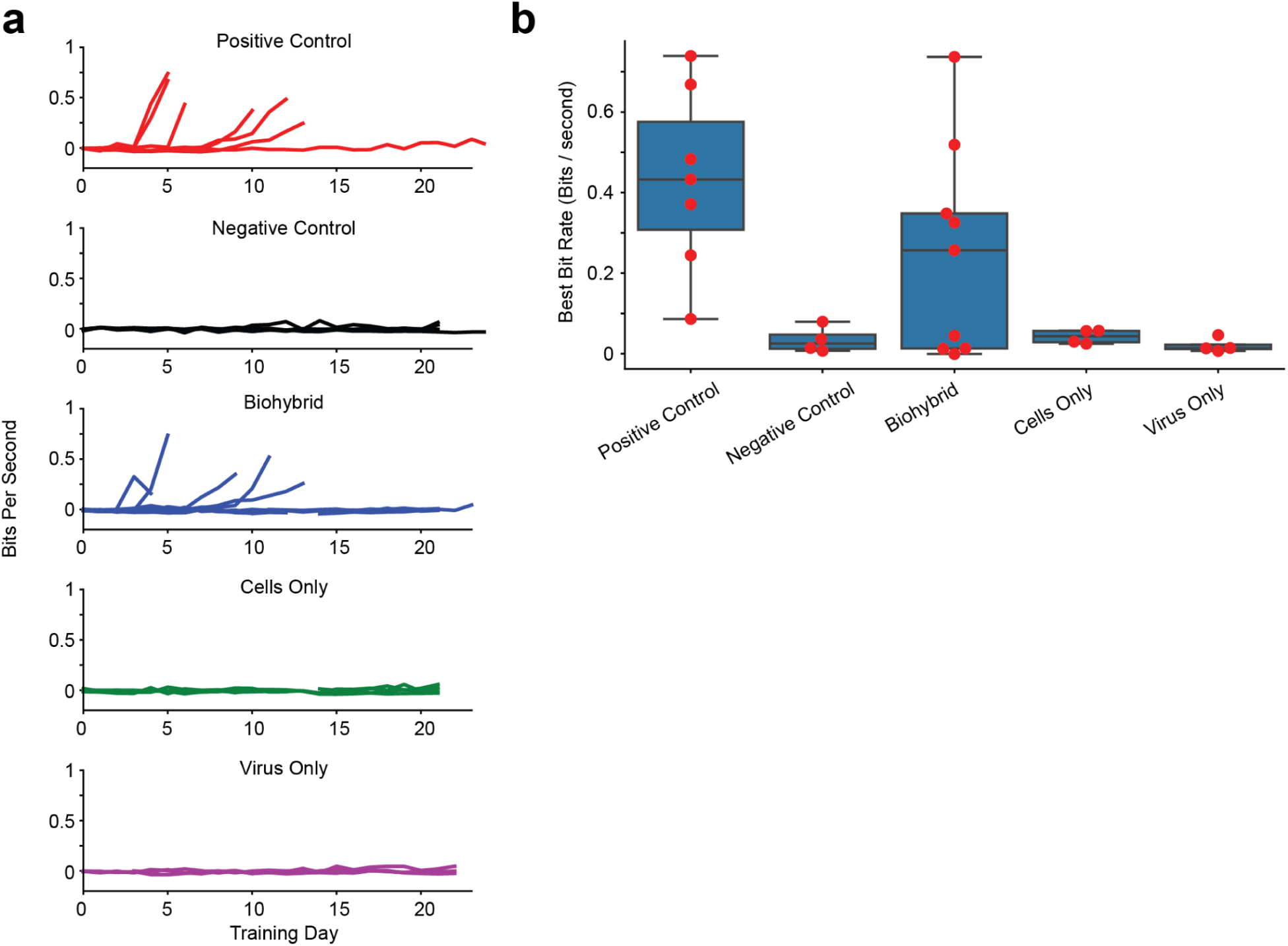
Positive bit rates using a biohybrid BCI. a. Plots showing training day (*x-*axis) versus mean bit-rate (bits per second) on the *y*-axis. Each colored line is the bit rate from an individual animal on a given training day. Subplot are positive control (AAV-excitatory opsin injected into cortex, n=7, red), negative control (jRGECO1a AAV injected into cortex, n=4, black), biohybrid implants (engrafted neurons on microwell scaffolds with CheRiff-eGFP, n=9, blue), cells only (microwell scaffolds loaded with cells and no AAV, n=4, green), or virus only (microwell scaffolds incubated with CheRiff-eGFP, n=4, magenta). b. Box and swarm plots showing the bit rate for the best session for each animal in each condition. Each dot corresponds to a single animal, and the boxplots show mean and interquartile range.

**Supplementary Figure 4:**
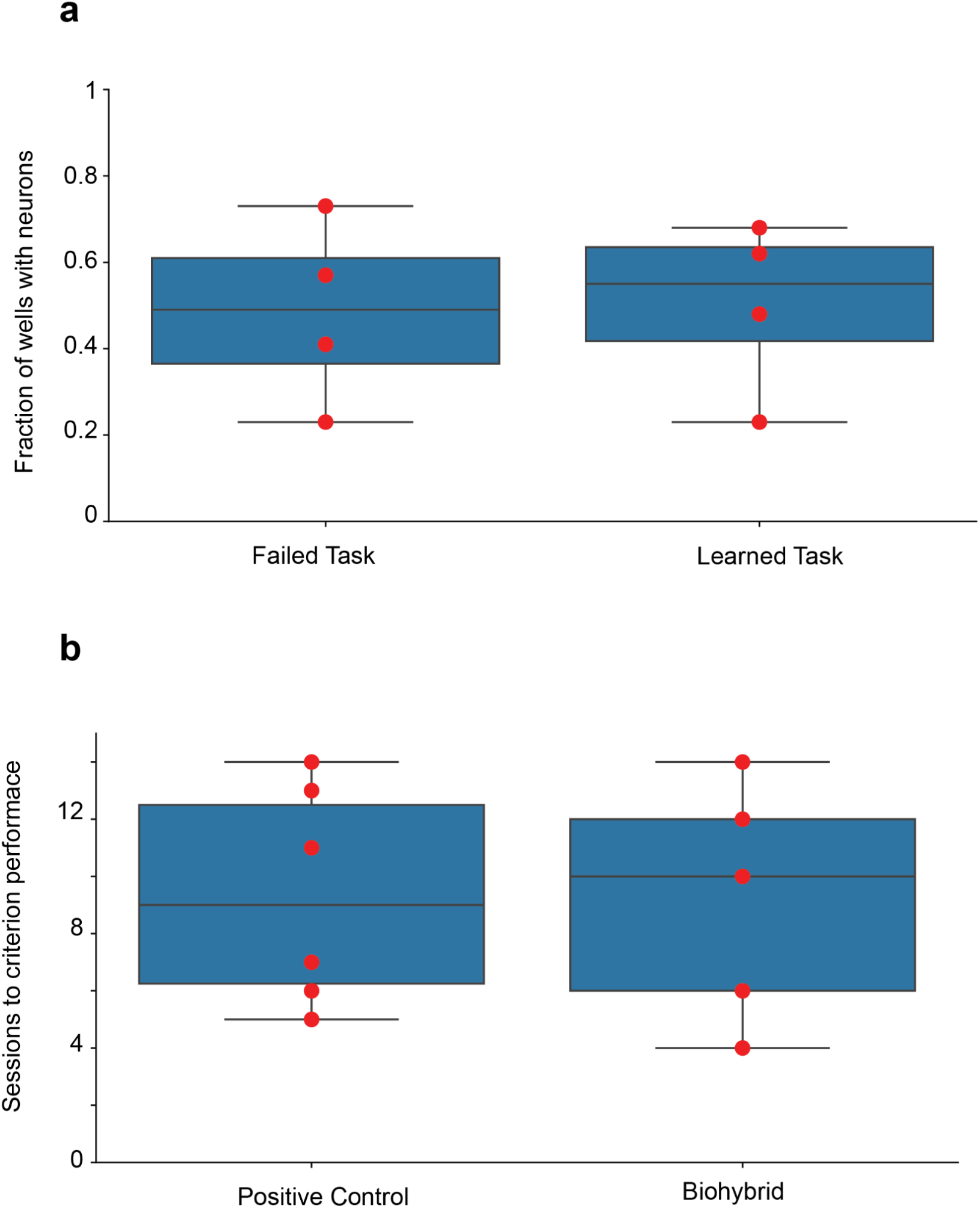
Time to learn optical stimulation task. a. Box and swarm plots showing the fraction of loaded microwells in the biohybrid animals that learned or did not learn the optical microstimulation task, each red dot corresponds to one animal (P>0.9, two-sided t-test). b. Box and swarm plots showing the number of training sessions needed to achieve criterion performance. Only animals that achieved criterion performance are shown, each red dot corresponds to one animal (P>0.95, two-sided t-test).

## Acknowledgements

We would like to acknowledge Emma Zhou for software support, Corey Wolin, and Bobby Urmy for hardware and facilities support, Elizabeth Carroll for insightful comments on the manuscript, and Rebecca King for animal husbandry support.

Work was performed in part in the nano@Stanford labs, which are supported by the National Science Foundation as part of the National Nanotechnology Coordinated Infrastructure under award ECCS-2026822

## Contributions

J.B. performed surgeries, built and ran the behavior assays, and conceptualized the project

P.D. performed single embryo dissections and cell culture.

E.Y. and S.S. built and tested cell loading fixtures

A.E.R. and M.E. provided imaging and histology support

A.R. helped train mice.

K.Z. and Y.K. designed and fabricated the cell scaffolds and conceptualized the project

M.H. conceptualized the project

A.M. conceptualized the project, performed 2P and confocal imaging and histology, wrote the manuscript and analyzed data.

